# Diversify your workflow! - An inconvenient suggestion to analyze spike data from intracranial recordings

**DOI:** 10.1101/2021.03.10.434718

**Authors:** Şükrü Okkesim, Shavika Rastogi, Olaf Christ, Peter Hubka, Nicole Rosskothen-Kuhl, Ulrich G. Hofmann

## Abstract

An important challenge of neuroscience research and future brain machine interfacing is the reliable assignment of spikes to putative neurons. By means of extracellular recordings, researchers try to match different types action potentials with their putative neuronal source and timing. Unfortunately, this procedure is by far not standardized and reliable, leading to many different suggestions and as many differing results. It appears that sharing of data is thus hampered by different processing pipelines in different labs, thus playing along the reproducibility crisis in neurosciences. To systematically shed light on this issue, we present preliminary results of several easy event detection schemes on one data set, meant to illustrate the inconsistencies arising from different processing pipelines already in its initial step. The results indicate that thresholding choices alter findings due to a lack of a ground truth for spike sorting. We suggest to increase reliability in findings by only accepting and further processing events accepted by more than one processing pipeline.

## 1. Introduction

The accurate detection of neuronal activity is a vital step for many neuroscientific studies involving electrophysiologic in vivo recordings [1]. As this step may very well become essential in a novel class of closed loop electro-ceutic devices [2], it is worth to spend a little time on its pitfalls and shortcomings. To finally utilize neuronal activity for medical purposes, researchers rightly focus on extracellular recordings accomplished by intracranially implanted micro-electrodes [3]. As long as they are positioned in the vicinity (up to 50μm [4]) to neurons, even single action potentials can be measured against a reference somewhere in the brain. A huge number of studies about the activity of neurons recorded extracellularly have given a wealth of information related to brain functions, neuronal networks and even individual characteristics of single neuron’s shape of spike. [5–6]. In recent years, improvements in microelectrode array manufacturing has achieved key advances and thus of a wide assortment of neural probe devices have enabled large-scale recording from groups of neurons in various brain areas and even over a long period of time [7–9].

Processing large scale recording requires *spike detection* as an initial step. Spike detection is the process of detecting interesting events in the stream of raw and noisy signals, which may then in a following step be assigned as tentative action potentials related to actual neurons and resulting in biologically interpretable spike trains *(spike sorting)* [10–13]. In the past decades, many approaches have been offered for the whole process towards spike train identification. However, seemingly each research group determines the set of parameters needed by its match with the purpose of their study depending on their historical experience. Thus, results for the same data set show human error and inconsistencies [14 – 15]. As a result, a method that has been shown to be successful in one lab may yield less successful results in another study [14 – 15]. Therefore, validating the reliability of spike analyzing methods is bound by ambiguities and prevents the establishment of a Gold-standard method minimizing ambiguity and enabling consistency and reliability.

As – according to the “garbage in-garbage out” rule of machine learning – the initial spike detection step is of critical importance.

Wide bandwidth raw data is commonly bandpass filtered between 300 Hz and 3 kHz to remove local field potentials (a source of information in its own right, 0.1 Hz to 300 Hz) and slow DC drift. Within this signal stream the interesting events may be identified by a wealth of methods but more often than not by plain vanilla thresholding.

Thresholding applies an amplitude threshold and determines interesting points/events passing this threshold, that is, the time stamped maxima of apparent spikes in the recorded waveform. Determining events by this method accepts a strong bias towards close-by or strongly firing neurons. Weaker signal sources may be hidden in the subthreshold noise level and thus are not thought to count towards the maximum detectable neurons. Worse still, choosing the right threshold is ambiguous in itself and adds bias to the selection, as there is a wide variability of signal characteristics, both intra- and inter-animals.

In this study, different spike detection methods by thresholding and their effects on the findings are discussed to reveal the inconsistencies of the result as a purpose.

## 2. Materials and Methods

Intracranial waveforms were recorded in anesthetized rats by a high density, 1600-electrode, 32-channel probe from multiple brain areas as presented in [7]. A needle-shaped shaft is divided into 50 blocks which each one of them consisting of 4 × 8 electrodes with an area of 17 × 17 μm^2^ (see Fig. 1).

**Fig. 1.**
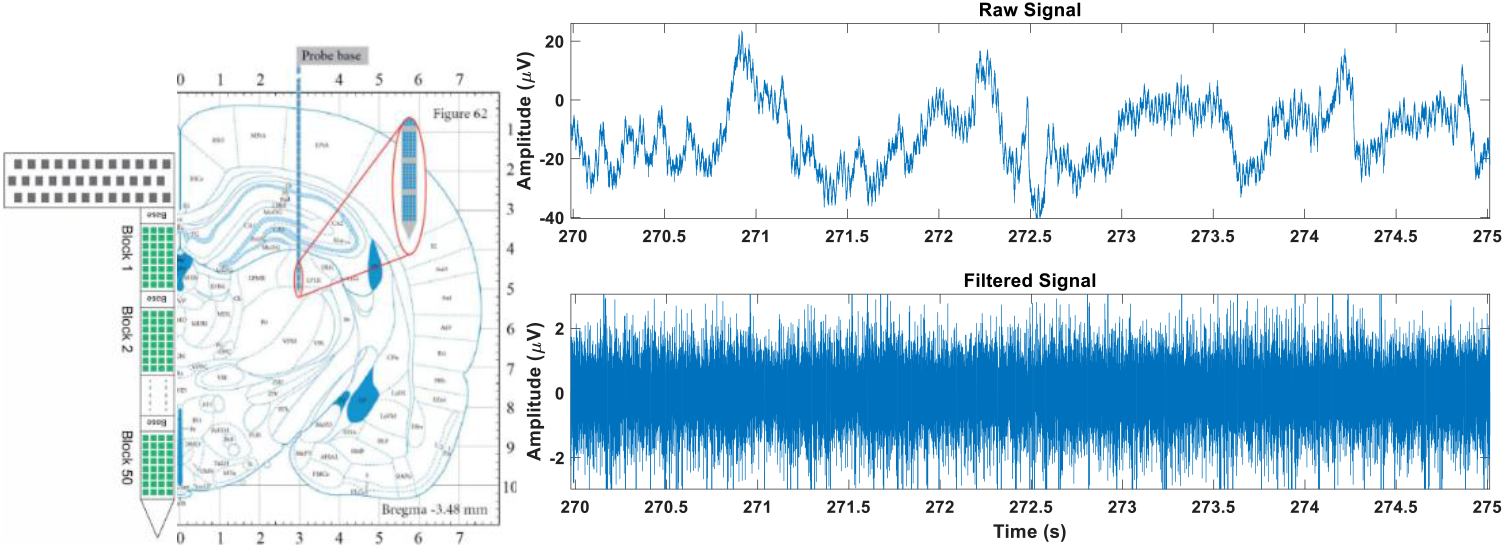
Position Image of 32-channel probe modified from [16]. with raw and filtered signal

Anesthetized Sprague Dawley **rats** (female, ca. 290 g) were used for in vivo experiments. All experimental protocols were approved by the Animal Care Committee of the University of Freiburg, Germany and the responsible Regierungspräsidium Freiburg (TVA licence G13/02). A stereotaxic frame from David Kopf Instruments (Tujunga, CA, USA) was used to immobilized the rats and enable precise final implantation of the probe. Coordinates of penetration were - 3.48 mm anterior-posterior, 3.0 mm medial-lateral and 5.0 mm dorsal-ventral into the lateral posterior thalamic nucleus according to Ref. [7, 16].

Recordings were sampled at 5 kHz by the wireless head stage while recording 32 channels (one block) at the same time for 30 seconds. A noncausal bandpass filter between 300 Hz and 2500 Hz was used (see Fig. 1.).

Though semiconductor technology allows the simultaneous recording of many adjacent sites, it increases the noise level due to small site’s huge impedance and flicker noise. Thus, already noisy neural signals provide even lower SNR [17].

**Wavelet denoising** is a poweful, yet easily tunable and common method, and there is evidence that the use of spike sorting with wavelet denoising leads to more successful neuroprosthetic systems [18]. In general, denoising is performed by converting the noisy data into an orthogonal time-frequency space - here by wavelets - and applying a threshold to the coefficients to remove the presumed noise in that space and lastly reconstruct it in in the original space. Like spike detection, performance of the wavelet denoising is influenced by several parameters like selection of mother wavelet, threshold selection ruler, decomposition level etc. [18 – 19]. However, the aim of the this study is not to discuss the inconsitency in the parameters selection procedure of wavelet denoising but of initial spike detection.Therefore, the parameters for wavelet denoising were selected as follows to be compatible with Ref [18]:

- Mother Wavelet: Symlet 7
- Decomposition Level: 3
- Threshold Rule: Hard

Another method to reveal the spike related peaks is the **nonlinear energy operator (NEO)** which is reported to be quite effective to detect peaks and edges in signals. To expose the peak of a spike, NEO produces an output relative to the product of the instantaneous amplitude and frequency of the input signal. As seen from the Eq1 for the NEO of a discrete-time sequence x(n), it is computationally simple and thus has an advantage to implement a real-time application [17 – 20].

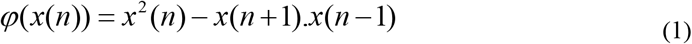

Inspired by the NEO another discrete-time sequence operator called as Shift-And-Multiply operator (SAM) was presented in [21]. Broad peaks are thought to be accentuated while shorter peaks caused by background noise are stifled by applying the SAM operator (Eq2).

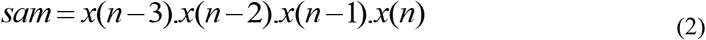

An important proposal towards a thresholding method with reduced ambiguity is offered by Quiroga et al. to solve an issue by very high threshold values for high firing rates and large spikes [1]. The presented threshold (Eq 3) is based on the median of the discrete-time sequence x(n) and is thought to diminish effect of interference of large spikes [15].

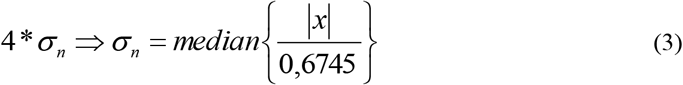

Alternatively, to the 4σ threshold, we implemented the Median Absolute Deviation (MAD) as presented in [22] (Eq 4).

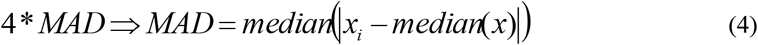

## 3. Results and Discussion

Within the past decade there has been much progress achieved in spike sorting especially in the recording technology of using multiple electrodes. Using threshold to detect events in huge data sets prior to spike sorting is a common step not only because of its simplicity but also because it is based on the amplitude which is the most evident feature of any spikes.

Figure 2 shows the exemplary, but no surprising results when applying a combination of above mentioned simple operators. Quite clearly, applying 4σ thresholding to the raw signal gives a quite a different spike count than MAD thresholding (Fig. 2, first row). Over a 30 sec stretch of the same data that is 57 spikes for 4σ vs. 1201 spikes for MAD (1.9 spikes/sec with 4σ vs. 40.03 spikes/sec with MAD). The discrepancy improves when the raw signal is denoised in a smart way by wavelet denoising and standard setting. Resulting spike counts over the same 30 sec is then 5470 for denoised-MAD thresholded data and 4475 for denoised-4σ thresholded data, respectively.

**Fig. 2.**
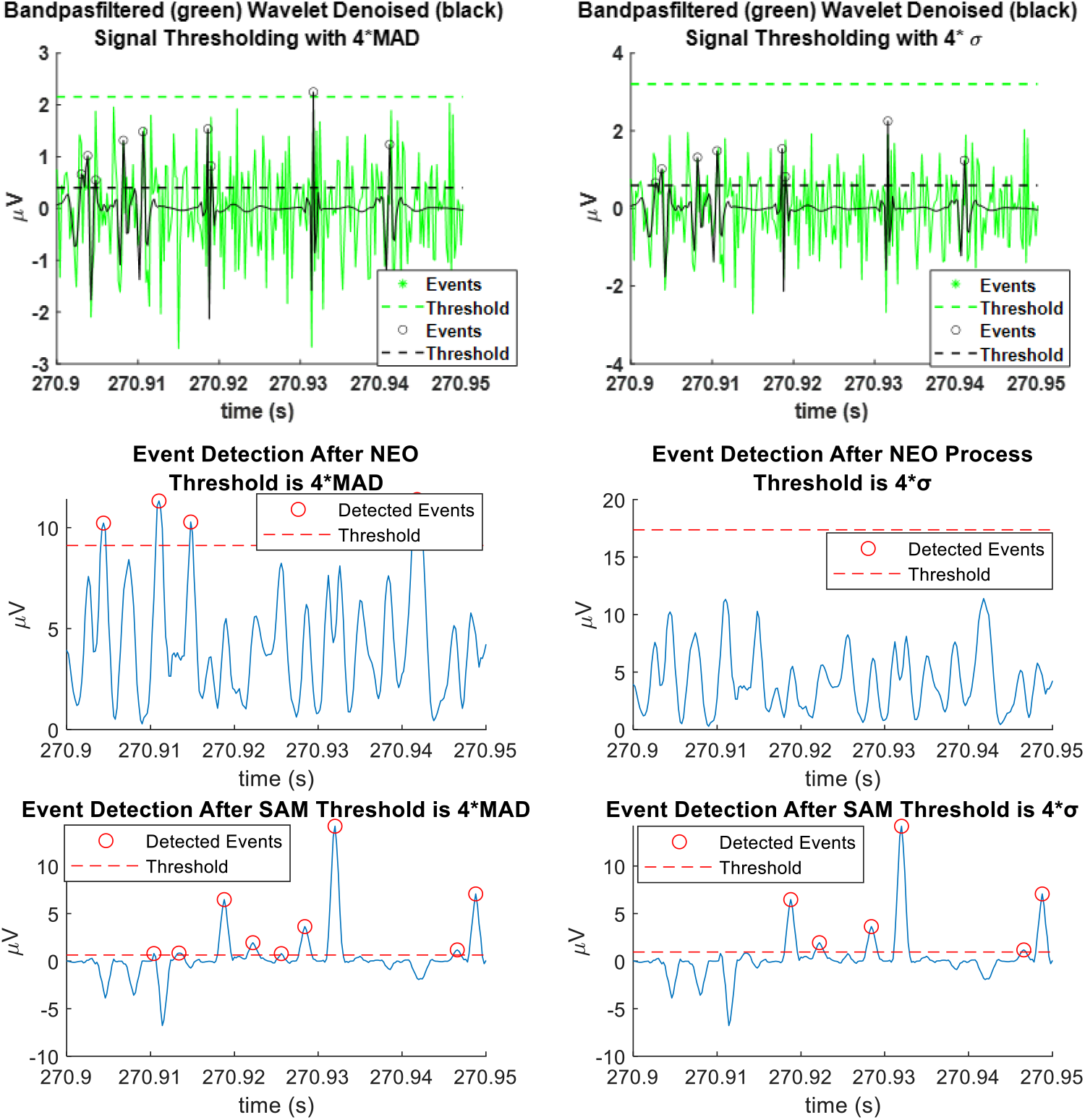
Detected events with two different thresholding

As thresholding is part of the event detection by NEO or SAM operators, their results do differ as well. NEO-processed, raw signals followed by MAD thresholding provide 98.83 spikes/sec, whereas NEO followed by 4σ leads to 20.97 spikes/sec. The values for SAM-processed raw signals are: 90.50 spikes/sec for 4σ thresholded and 119.83 spikes/sec for MAD-thresholded data. Table 1 presents these results in a more accessible way.

**Table 1.**
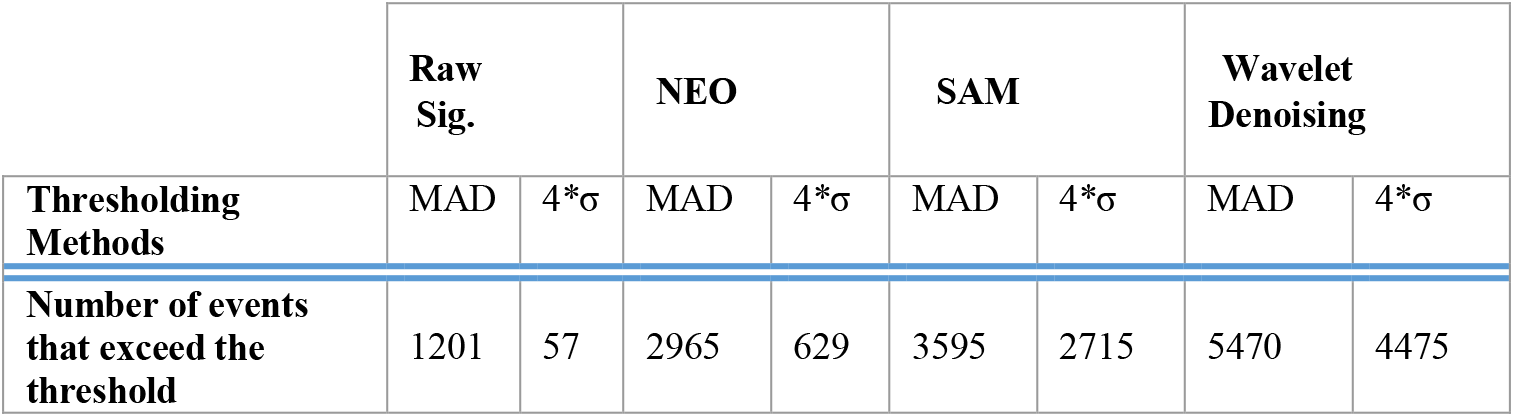
Number of the peaks that cross the Threshold for 30 sec signal

We are well aware, that a more thorough analysis of all our data sets across all measured animals and locations is needed (and under way). However, the presented results already cast doubt on the robustness and reliability of some current spike detecting schemes implemented in silico. This in turn has severe consequences for their further use in spike sorting (for neuroprosthetic use) and spike train analysis (like tuning curve analysis). As automated processing pipelines in future BMI are not easily updated by novel, better suitable analysis algorithms while implanted, the methods to be deployed need to be as reliable and robust as possible.

In addition, as there is no true ground truth data available while doing in vivo recordings, there is no way of truly benchmarking detection schemes in situ, as the recording situation most likely does not stay stationary.

In order to improve both aspects, reliability and robustness, we propose a brute force solution: Only spikes detected with more than one method are worth further considering. That way at least a subset of detected events gain a high reliability and may be used for further use like control signals in BMIs. Figure 3 shows a Venn graph for each of the thresholding methods intended to clarify the resulting spike counts common to the different peak points.

**Fig. 3.**
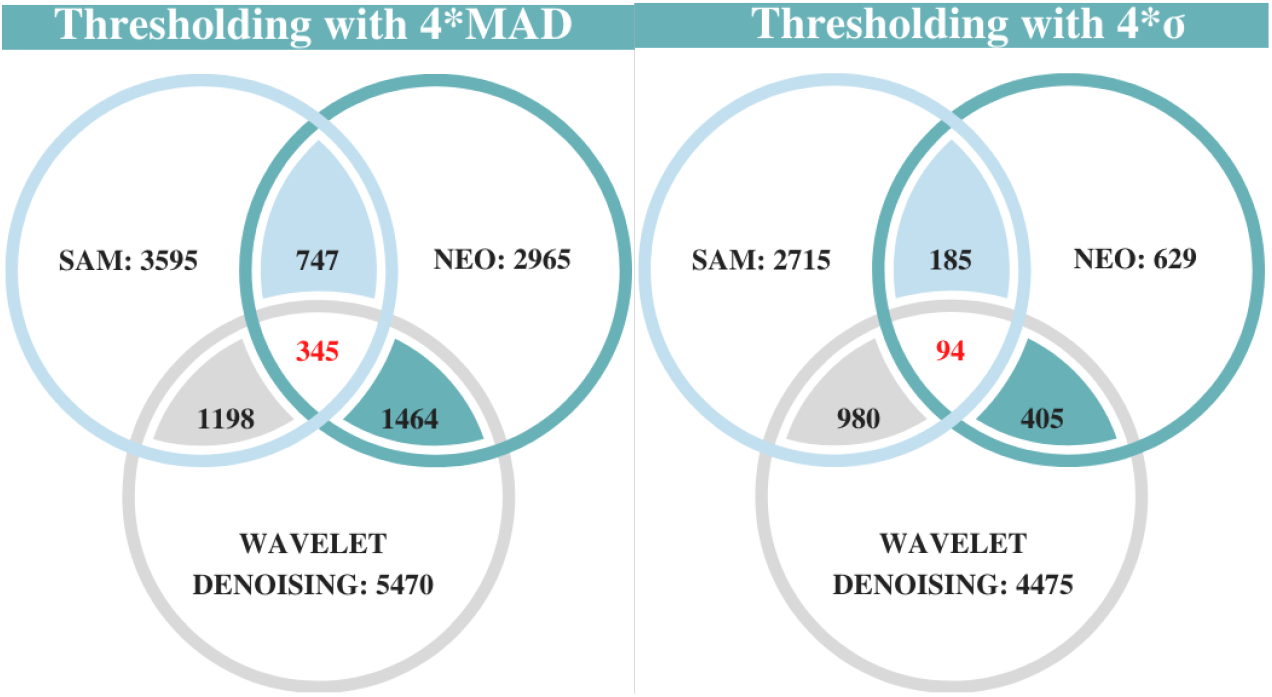
Venn Diagram of the events that cross the Threshold

Ongoing work on the full data set at our disposal and data sets from different other electrophysiologic experiments will hopefully shed light on the question, whether or not this strong confinement to multiple-assured spikes will help robustness of analyzes and use of neuronal spike data.

## Acknowledgments

Part of this work was supported by Alexander von Humboldt Foundation-Germany, through The Philipp Schwartz Initiative grant to one of the authors.

## References

1. Quian Quiroga R, Nadasdy Z, Ben-Shaul Y Unsupervised Spike Detection and Sorting with Wavelets and Superparamagnetic Clustering. Neural Comp 16:1661–1687, (2004).

2. Fanselow EE., Reid AP., Nicolelis MA.: Reduction of pentylenetetrazole-induced seizure activity in awake rats by seizure-triggered trigeminal nerve stimulation. J Neurosci. 20(21), 8160–8168, (2000).

3. Ajiboye, A.B., F.R. Willett, D.R. Young, W.D. Memberg, et al: Restoration of reaching and grasping movements through brain-controlled muscle stimulation in a person with tetraplegia: a proof-of-concept demonstration. The Lancet. 389(10081): 1821-1830, (2017).

4. Buzsáki G. Large-scale recording of neuronal ensembles. Nat Neurosci. 7(5):446–451, 2004.

5. Rey H. G., Pedreira C., Quiroga R. Q.: Past, present and future of spike sorting techniques. Brain Research Bulletin 119 (B), 106–117 (2015)

6. Lewicki, Michael S.. A review of methods for spike sorting: the detection and classification of neural action potentials. Network 9 (4), R53–78 (1998).

7. A. Sayed Herbawi et al., CMOS Neural Probe With 1600 Close-Packed Recording Sites and 32 Analog Output Channels. Journal of Microelectromechanical Systems, vol. 27(6), 1023–1034, (2018).

8. Luan, L., X. Wei, Z. Zhao, J.J. Siegel, et al: Ultraflexible nanoelectronic probes form reliable, glial scar–free neural integration. Science Advances. 3: p. e1601966 (2017).

9. Böhler, C., C. Kleber, N. Martini, Y. Xie, I. Dryg, T. Stieglitz, U.G. Hofmann, and M. Asplund, Actively controlled release of Dexamethasone from neural microelectrodes in a chronic in vivo study. Biomaterials. 129: p. 176–187 (2017).

10. Souza, B.C., V. Lopes-Dos-Santos, J. Bacelo, and A.B.L. Tort, Spike sorting with Gaussian mixture models. Sci Rep,. 9(1): 3627 (2019).

11. Chung, J.E., J.F. Magland, A.H. Barnett, V.M. Tolosa, et al A Fully Automated Approach to Spike Sorting. Neuron,. 95(6): p. 1381–1394 (2017).

12. Luan, S., I. Williams, M. Maslik, Y. Liu, F. et al, Compact standalone platform for neural recording with real-time spike sorting and data logging. J Neural Eng,. 15(4): p. 046014 (2018).

13. Quiroga R. Q. Spike sorting, Scholarpedia, 2(12):3583, (2007).

14. Sukiban J., Voges N., Dembek T.A., et al., Evaluation of Spike Sorting Algorithms: Application to Human Subthalamic Nucleus Recordings and Simulations. Neuroscience, 414:168–185 (2019)

15. Menne, K. & Hofmann, U. Automatic neurophysiological identification of the human subthalamic nucleus for the implantation of deep brain stimulation electrodes based on statistical signal processing methods and unsupervised classification. 3rd European Medical & Biological Engineering Conference 2005, vol. 1627 pp.824–829, (2005).

16. Paxinos G. and Watson C., The Rat Brain in Stereotaxic Coordinates. Elsevier, Amsterdam, The Netherlands (2005).

17. Kim, K.H. and S.J. Kim, Neural spike sorting under nearly 0-dB signal-to-noise-ratio unsing nonlinear energy operator and artificial neural-network classifier. IEEE Trans. Biomed. Eng,. 47: p. 1406–1411, (2000).

18. Citi L, Carpaneto J, Yoshida K, et al. On the use of wavelet denoising and spike sorting techniques to process electroneurographic signals recorded using intraneural electrodes. J Neurosci Methods, 172(2): 294–302 (2008).

19. Emeterio J.L.S., Pardo E., Rodríguez M.A. (2011) Denoising Ultrasound RF Signals by Wavelet Cycle Spinning Shrinkage. In: Jobbágy Á. (eds) 5th European Conference of the International Federation for Medical and Biological Engineering. IFMBE Proceedings, vol 37, pp.78–81. Springer, Berlin, Heidelberg, (2011).

20. Mukhopadhyay S. and Ray G. C., A new interpretation of nonlinear energy operator and its efficacy in spike detection, IEEE Trans. Biomed. Eng., vol. 45, 180–187, (1998).

21. T. Gritsun et al., A Simple Microelectrode Bundle for Deep Brain Recordings, 2007 3rd International IEEE/EMBS Conference on Neural Engineering, Kohala Coast, HI, pp. 114–117 (2007).

22. James J. Jun, Catalin Mitelut, Chongxi Lai, et al., Real-time spike sorting platform for high-density extracellular probes with ground-truth validation and drift correction. bioRxiv., 10.1101/101030, (2017)

